# Plasma-activated water hydrogel enables effective clearance of wound-associated microbial infections

**DOI:** 10.1101/2025.11.10.687761

**Authors:** Debapriya Mukherjee, Pallab Ghosh, Raju S Rajmani, Kavita Verma, Akhil Gowda, Apoorva Birua, Lakshminarayana Rao, Dipshikha Chakravortty

**Author notes:** Corresponding Authors Lakshminarayana Rao, Centre for Sustainable Technologies, Indian Institute of Science, Bangalore, Karnataka 560012, India., Dipshikha Chakravortty, Department of Microbiology and Cell Biology, Indian Institute of Science, Bangalore, Karnataka 560012, India. Equal contribution (Debapriya Mukherjee, Pallab Ghosh).

## Abstract

Surgical site infections are among the most common postoperative complications and remain a major global health concern, further intensified by increasing antimicrobial resistance. Plasma-activated water (PAW) has emerged as a novel antimicrobial strategy, though earlier studies suggested its activity depends on an acidic pH. To overcome this limitation, we previously developed a high-strength, neutral pH–buffered formulation (hs-PAbW) optimized for human clinical use, which preserved strong antimicrobial properties. In the present work, we created a hydrogel incorporating hs-PAbW at a bactericidal yet non-cytotoxic concentration (hs-PAbW-20) and evaluated its therapeutic performance in mouse wound models. In Balb/c mice, hs-PAbW-20 gel markedly reduced *Pseudomonas aeruginosa* and *Staphylococcus aureus* loads, demonstrating efficacy comparable to neomycin and soframycin. The gel also enhanced wound closure by regulating key genes involved in tissue repair. In a diabetic C57BL/6 model, hs-PAbW-20 gel similarly lowered *P. aeruginosa* burden and accelerated healing. Collectively, these results highlight hs-PAbW-20 gel as a promising next-generation wound therapy capable of addressing the shortcomings of traditional antibiotic-based treatments.

## Introduction

Surgical site infections (SSIs) rank among the most common postoperative complications and represent a major global health burden (1). According to CDC NNIS data, SSIs constitute 14–16% of all hospital-acquired infections and 38% in surgical patients, significantly increasing morbidity and mortality (2–4).

SSIs are predominantly caused by *Staphylococcus aureus*, *Pseudomonas aeruginosa*, *Klebsiella pneumoniae*, and *Acinetobacter baumannii* (5–8). Escalating antimicrobial resistance severely limits conventional therapies. An emerging antimicrobial approach is plasma-activated water (PAW), generated by exposing water to cold plasma, an ionized gas containing electrons, ions, and neutral atoms (11–13). Plasma–water interactions produce reactive oxygen and nitrogen species, which confer antimicrobial activity (14–16). Traditional PAW requires acidic pH for efficacy, but we previously developed a high-strength, neutral-pH bicarbonate-buffered version (hs-PAbW) that retains strong bactericidal activity against ESKAPE pathogens via RNS-mediated peroxynitrite and nitrotyrosine formation (17–20).

Here, we formulated hs-PAbW into a non-cytotoxic hydrogel (hs-PAbW-20-gel) and evaluated its therapeutic potential in murine wound-infection models. The hydrogel effectively eradicated *P. aeruginosa* and *S. aureus*, accelerated wound closure, and matched or surpassed the performance of standard topical antibiotics. These findings position hs-PAbW-20-gel as a promising alternative for managing infected wounds.

## Materials and methods

### Generation of hs-PAbW-20-gel

hs-PAbW was prepared by treating 50 mL of 25 mM bicarbonate water with a 10 kV, 20 kHz AC plasma discharge (aluminium electrodes, 5 mm gap, 1 L/min air flow, 32 ± 2 W) in a 100ml glass reactor for 60min (repeated ≥3 times) (17, 19, 20). The resulting hs-PAbW was diluted 1:5 with 25mM bicarbonate water to obtain a 20% solution, to which 0.5%(w/v) food-grade xanthan gum was added to form the hs-PAbW-20-gel.

### Bacterial strains and culture conditions

Methicillin-resistant *Staphylococcus aureus* (Isolate-SA3) was obtained from Prof. Pradyut Prakash (Banaras Hindu University), and *Pseudomonas aeruginosa* PAO1 from Prof. Anne-Béatrice Blanc-Potard (Université de Montpellier). The bacteria were cultured overnight in LB broth at 37°C with 170rpm shaking to stationary phase, then diluted 1:100 and cultured for 6h under the same conditions to obtain log-phase cells.

### Propidium Iodide staining

Log-phase *P. aeruginosa* and *S. aureus* (10⁸ CFU) were treated with 100 µl hs-PAbW-20-gel or control gel and incubated overnight at 37°C. Cells were stained with 1 µg/mL propidium iodide (PI) (Sigma), and cell death was assessed by flow cytometry (BD FACSVerse) using FACSuite software (20–22).

### Eukaryotic cell lines and growth conditions

L929 mouse fibroblasts (ATCC) were cultured in DMEM(Sigma-Aldrich) supplemented with 10%FCS (Gibco) and maintained at 37°C in a 5%CO₂ humidified incubator.

### *In-vitro* cell migration assay

1×10⁵ cells/well were seeded in 24-well plates and cultured overnight to confluence. Monolayers were washed with 1×PBS, scratched with a sterile 200µl pipette tip, and washed again to remove debris. Cells were treated with hs-PAbW-20-gel or control gel (1:5 v/v in culture medium) and incubated for 24h. Scratch images were captured at 0h and 24h using a microscope-mounted mobile camera, and closure was quantified by measuring scratch width with ImageJ (23, 24).

### Ethics statement for animal experiments

All animal experiments were approved by the Institutional Animal Ethics Committee (IAEC), IISc Bangalore (Reg. No. 48/1999/CPCSEA; Approval No. CAF/Ethics/973/2023) and performed in strict compliance with CPCSEA guidelines.

### Wound healing experiment in mice

Wound healing was assessed in 10–12-week-old Balb/c and C57BL/6 mice using a modified excisional wound model (25). Mice were anesthetized with ketamine (120 mg/kg, i.v.), dorsally depilated, and two full-thickness 10-mm wounds were created with a sterile biopsy punch. Wounds were inoculated with 10⁵ CFU *P. aeruginosa* or 10⁴ CFU *S. aureus* and allowed to establish infection for 30min. Starting on 0d, ∼100 µl of hs-PAbW-20-gel, control gel, or commercial antibiotic cream (neomycin or soframycin) were applied topically every alternate day until 7d. Wounds were photographed, and mice were euthanized on 9d for tissue collection. Wound diameter was quantified using ImageJ.

### Generation of diabetic mice model

Type 1 diabetes was established in 10–12-week-old male C57BL/6 mice using streptozotocin via five consecutive 40 mg/kg intraperitoneal injections (26). During this induction, mice were provided 10% sucrose water, which was replaced with regular drinking water from 6d onward, while they were maintained on a standard chow diet and monitored for blood glucose levels using a Dr. Morepen BG-03 glucometer. Seven days post-STZ induction, diabetic mice received a full-thickness excisional wound that was subsequently infected by inoculation with 10^4^ CFU of *P. aeruginosa* before undergoing specified treatments.

### Bacterial burden enumeration by spread plating

At 9dpi, wound tissues were harvested in 1×PBS, homogenized by bead beating, serially diluted, and plated on selective agar (cetrimide agar for *P. aeruginosa*; mannitol salt agar for *S. aureus*). Following overnight incubation at 37°C, CFU were enumerated, corrected for dilution, and normalized to tissue weight to determine bacterial burden (CFU/gm-weight).

### RNA Isolation and RT-qPCR

At 9dpi, wound tissues were collected in TRIzol (RNAiso Plus, TaKaRa) and stored at −80°C. Total RNA was extracted by phenol–chloroform, DNase-treated (Invitrogen TURBO DNase, AM2239), and reverse-transcribed using the RevertAid First Strand cDNA Synthesis Kit (Thermo Fisher K1622). RT-qPCR was performed with PowerUp SYBR Green Master Mix (A25742) on a QuantStudio 5 system. Gene expression was normalized to β-actin.

### Histopathology imaging

At 9dpi, wound tissues were fixed in 3.5% paraformaldehyde and processed for histology with haematoxylin–eosin staining. Images were acquired using light microscopy and evaluated by a qualified veterinary pathologist.

### Statistical Analysis

Biological and technical replicates are detailed in figure legends. Statistical tests indicated per figure. Significance was set at *p*<0.05. Data are presented as mean±SD or mean±SEM as specified in legends. All analyses and graphs were performed using GraphPad Prism 8.4.3.

## Results

### hs-PAbW-20-gel maintained its antibacterial effectiveness without affecting cell migration

hs-PAbW-20-gel was formulated by diluting the hs-PAbW to a 20% concentration and solidified with xanthan gum, a non-cytotoxic ratio shown to retain anti-*S. aureus* activity (20). This gel formulation retained its bactericidal activity against chronic wound pathogens *P. aeruginosa* and *S. aureus* (**Figure 1A-C**) (27).

**Figure 1.**
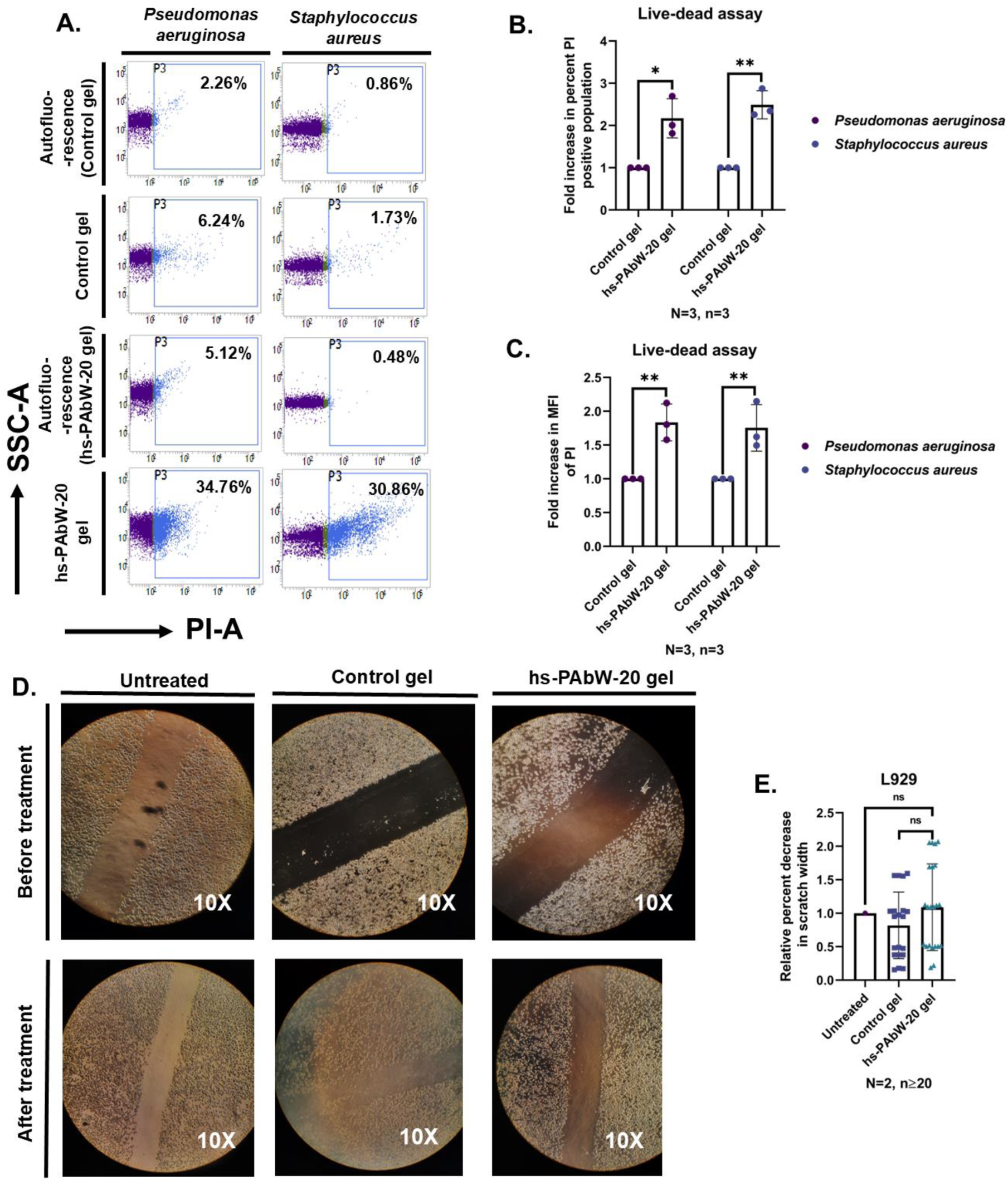
hs-PAbW-20-gel preserves antibacterial activity without impacting cell migration. **(A)** Representative flow cytometry profiles (SSC-A vs. PI-A) showing PI uptake in *P. aeruginosa* and *S. aureus* after overnight treatment with hs-PAbW-20 gel or control. **(B)** Fold increase in PI-positive cell populations. **(C)** Fold increase in PI MFI. **(D)** Representative images of L929 scratch assay at 0h and 24h post-treatment with hs-PAbW-20-gel or control. **(E)** Percentage reduction in scratch width after 24 h. Data in (B, C, E) are mean±SD. (A–C): N=3 biological replicates, n=3 technical replicates each; (D–E): N=2, n≥20 fields. Statistical tests: one-way ANOVA (B, C, E); ****p<0.0001, ***p<0.001, **p<0.01, *p<0.05.

Cell migration, critical to wound healing and other processes (28), was evaluated using scratch assay in L929 cells. The results demonstrated that hs-PAbW-20-gel treatment did not impair cell migration, with 24h closure rates being comparable to the control (**Figure 1D-E**). Collectively, these *in vitro* results demonstrate that hs-PAbW-20-gel retains bactericidal activity against *P. aeruginosa* and *S. aureus* while remaining non-cytotoxic and without impacting cell migration.

### hs-PAbW-20-gel reduced *P. aeruginosa* burden in wounds, with faster closure and elevated expression of pro-healing genes

The topical efficacy of the hs-PAbW-20-gel was evaluated in bilateral full-thickness wounds infected with *P. aeruginosa* across two distinct mouse strains. In Balb/c mice, the gel significantly accelerated wound closure **(Figure 2A–B**) and led to a reduction in bacterial load compared to control treatments (**Figure 2C**).

**Figure 2.**
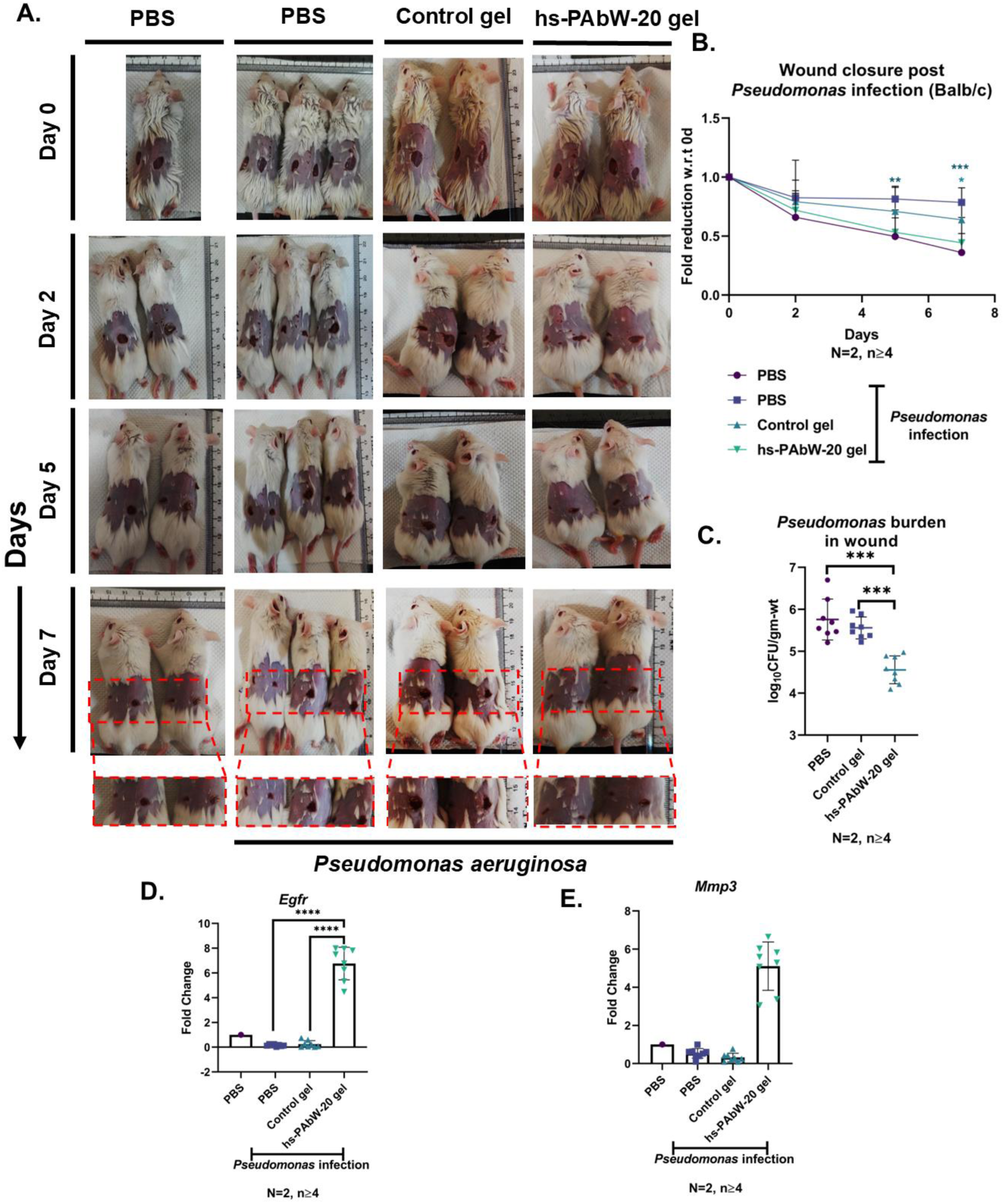
hs-PAbW-20-gel accelerates wound closure and reduces *P. aeruginosa* burden in infected Balb/c mice while upregulating healing-associated genes. **(A)** Representative images of *P. aeruginosa*-infected wounds under different treatment conditions. **(B)** Fold reduction in wound diameter relative to 0d. **(C)** *P. aeruginosa* burden in tissue at 9d post-infection. **(D–E)** Relative mRNA expression of (D) *Egfr* and (E) *Mmp3* in uninfected and infected wounds, treated as indicated. Data in (B–E) are mean±SD, pooled from N=2 independent experiments, n≥4 mice/group. Statistical tests: one-way ANOVA (B, D, E), two-way ANOVA (grouped data in B), Mann–Whitney U-test (C); ****p<0.0001, ***p<0.001, **p<0.01, *p<0.05.

Previous studies have highlighted the roles of matrix metalloproteinases (MMPs), epidermal growth factors (EGF), and platelet-derived growth factor (PDGF) in wound healing (29–31). We found that *Egfr* and *Mmp3* were significantly upregulated in hs-PAbW-20-gel–treated infected wounds relative to all controls (**Figure 2D–E**).

Validation in the C57BL/6 strain confirmed the ability of hs-PAbW-20-gel to clear *P. aeruginosa* infection (**Figure S1C**) and enhance the expression of *Pdgf*, *Egfr*, *Mmp1*, and *Mmp3* (**Figure S2A-D**). However, enhanced wound closure was not observed in the C57BL/6 model, possibly due to that strain’s characteristically Th1-biased, proinflammatory response (**Figure S1A-B**) (32). These results collectively demonstrate that the hs-PAbW-20-gel effectively mitigates infection and actively stimulates the transcriptional pathways essential for wound repair across different genetic backgrounds.

### hs-PAbW-20-gel demonstrated efficacy comparable to commercially available antibiotic creams

The efficacy of hs-PAbW-20-gel was directly benchmarked against standard antibiotic ointments (neomycin and soframycin) in Balb/c mice with full-thickness *P. aeruginosa* infected wounds (**Figure 3**). Compared to the control gel and both antibiotics, the hs-PAbW-20-gel demonstrated superior performance in accelerating wound closure by 7dpi, while effectively reducing the bacterial burden to levels comparable with both antibiotic treatments (**Figure 3A-C**). Upregulation of *Pdgf, Egfr,* and *Mmp1* was observed, mirroring the expression pattern observed with soframycin (**Figure S3A-D**).

**Figure 3.**
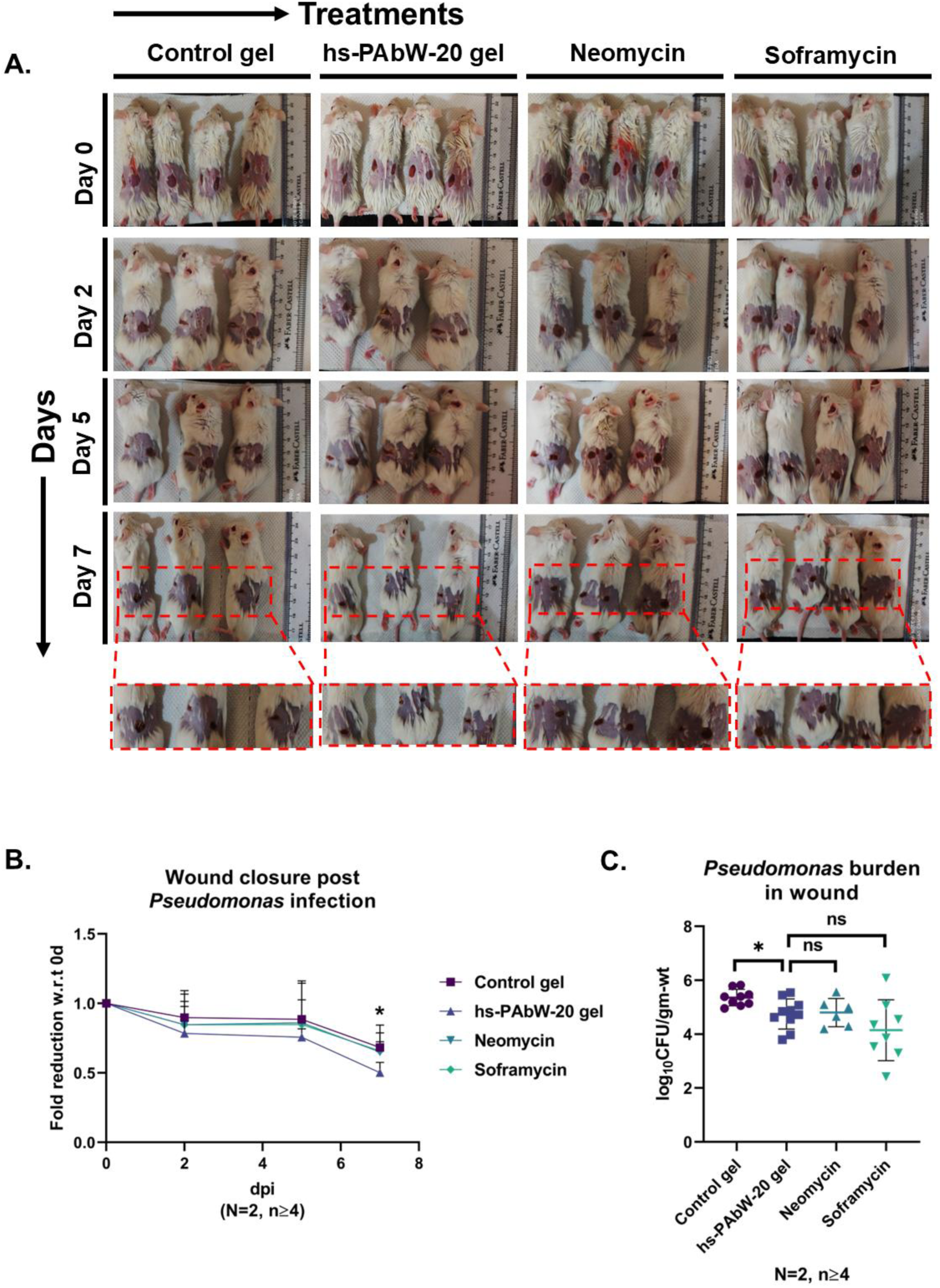
hs-PAbW-20-gel outperforms commercial antibiotic creams in promoting wound closure while matching their antimicrobial efficacy against *P. aeruginosa* in Balb/c mice. **(A)** Representative wound images (0d–7d) after *P. aeruginosa* infection and treatment with control gel, hs-PAbW-20-gel, neomycin, or soframycin. **(B)** Fold reduction in wound diameter relative to 0d. **(C)** *P. aeruginosa* burden in tissue at 9d. Data in (B–C) are mean±SD, pooled from N=2 independent experiments, n≥4 mice/group. Statistical tests: two-way ANOVA (B), one-way ANOVA (C, grouped data), Mann–Whitney U-test (CFU); ****p<0.0001, ***p<0.001, **p<0.01, *p<0.05.

Histological analysis confirmed the gel’s efficacy, as hs-PAbW-20-gel-treated wounds showed significantly improved tissue organization (score-2) and reduced inflammatory pathology compared to the severe pathology seen in control-treated wounds (score-4) (**Figure 4**).

**Figure 4.**
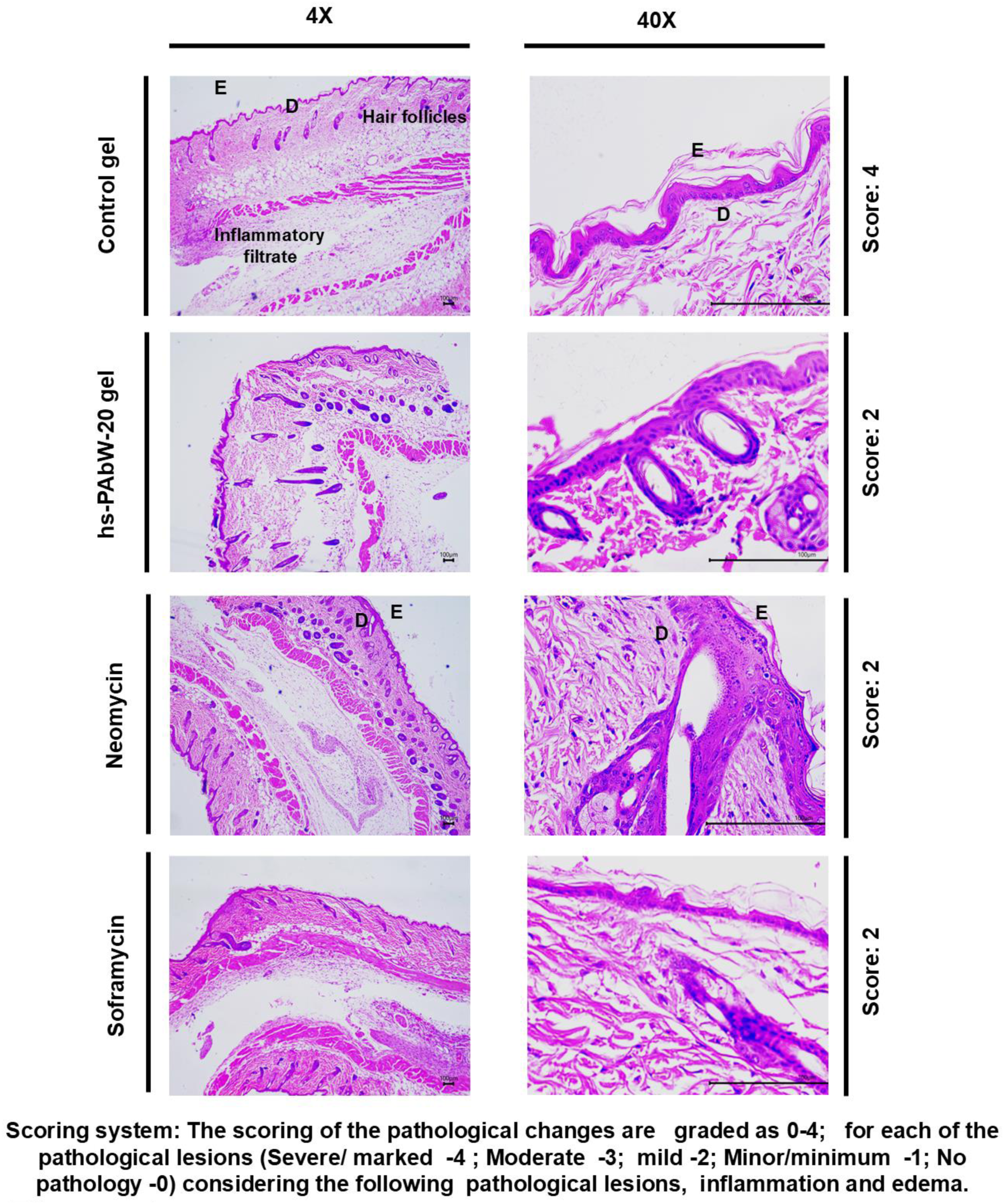
Histopathological evaluation of *P. aeruginosa*-infected wounds at 9d. H&E-stained sections showing tissue repair and epithelialization in wounds under treatment conditions.

### hs-PAbW-20-gel reduced *S. aureus* load in infected wounds, promoting accelerated closure like commercial antibiotic ointments

In experiments using Balb/c mice wounds infected with *S. aureus*, hs-PAbW-20-gel accelerated wound closure compared to control gel, performing comparably to neomycin and soframycin (**Figure S4A–B**). Furthermore, hs-PAbW-20 markedly reduced the *S. aureus* burden to levels like those achieved by neomycin (**Figure S4C**). The gel also upregulated *Pdgf, Egfr,* and *Mmp1* to an extent comparable to the antibiotic treatments (**Figure S5A–C**).

Histological analysis, while showing that the antibiotics achieved superior tissue pathology scores (0-1), revealed that the hs-PAbW-20-gel significantly improved the pathology score from 4 to 3 over the control gel, demonstrating its ability to reduce inflammation and promote tissue organization (**Figure S6**). Overall, the hs-PAbW-20-gel was also effective in clearing *S. aureus* infection, promoting wound healing, and exhibiting efficacy broadly equivalent to standard antibiotic creams.

### hs-PAbW-20-gel effectively reduces *P. aeruginosa* burden from infected wounds of diabetic mice, along with improved wound closure

To address the challenges of delayed healing and persistent infection in diabetic wounds (33–35), hs-PAbW-20-gel was tested in the streptozotocin-induced Type 1 diabetic C57BL/6 mice (**Figure S7A**) (36). By 7d, hs-PAbW-20-gel significantly accelerated wound closure compared to control gel (**Figure 5A–B**) and reduced *P. aeruginosa* better than antibiotic ointments (**Figure 5C**). *Pdgf*, *Egfr*, *Mmp1*, and *Mmp3* was strongly upregulated in hs-PAbW-20–treated wounds (**Figure S7B–E**).

**Figure 5.**
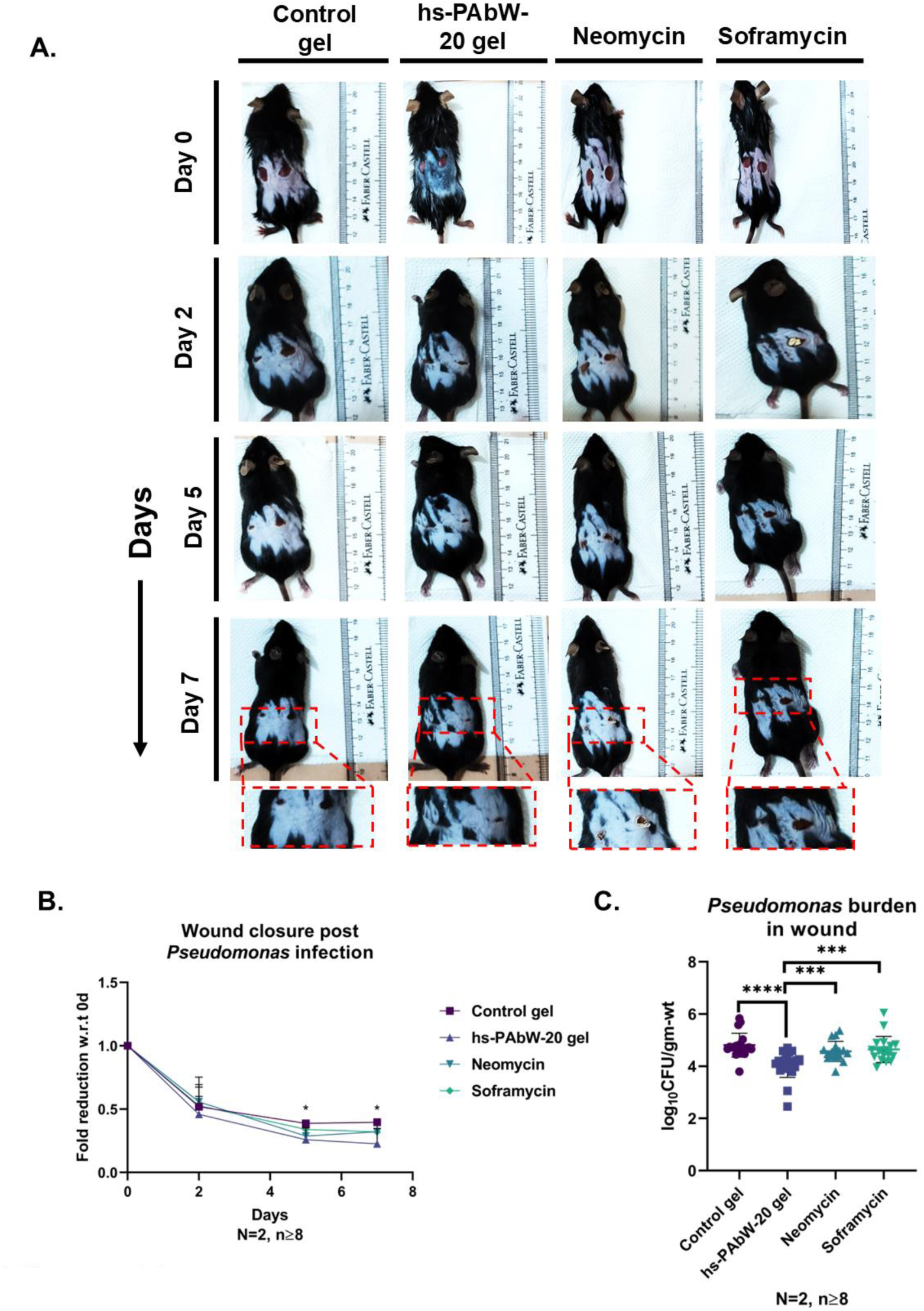
hs-PAbW-20-gel reduces *P. aeruginosa* burden and accelerates wound closure in diabetic C57BL/6 mice. **(A)** Representative wound images (0d–7d) after *P. aeruginosa* infection and treatment with control gel, hs-PAbW-20 gel, neomycin, or soframycin. **(B)** Fold reduction in wound diameter relative to 0d. **(C)** *P. aeruginosa* burden in tissue at 9d. Data in (B–C) are mean±SD, pooled from N=2 independent experiments, n≥8 mice/group. Statistical tests: two-way ANOVA (B), one-way ANOVA (C, grouped data), Mann–Whitney U-test (CFU); ****p<0.0001, ***p<0.001, **p<0.01, *p<0.05.

Histology revealed severe pathology (score-4) in control- and antibiotic-treated wounds, with necrosis and minimal re-epithelialization. However, hs-PAbW-20–treated wounds showed marked improvement (score-2), including restored epithelium, keratinization, angiogenesis, and early hair follicle regeneration (**Figure 6**).

**Figure 6.**
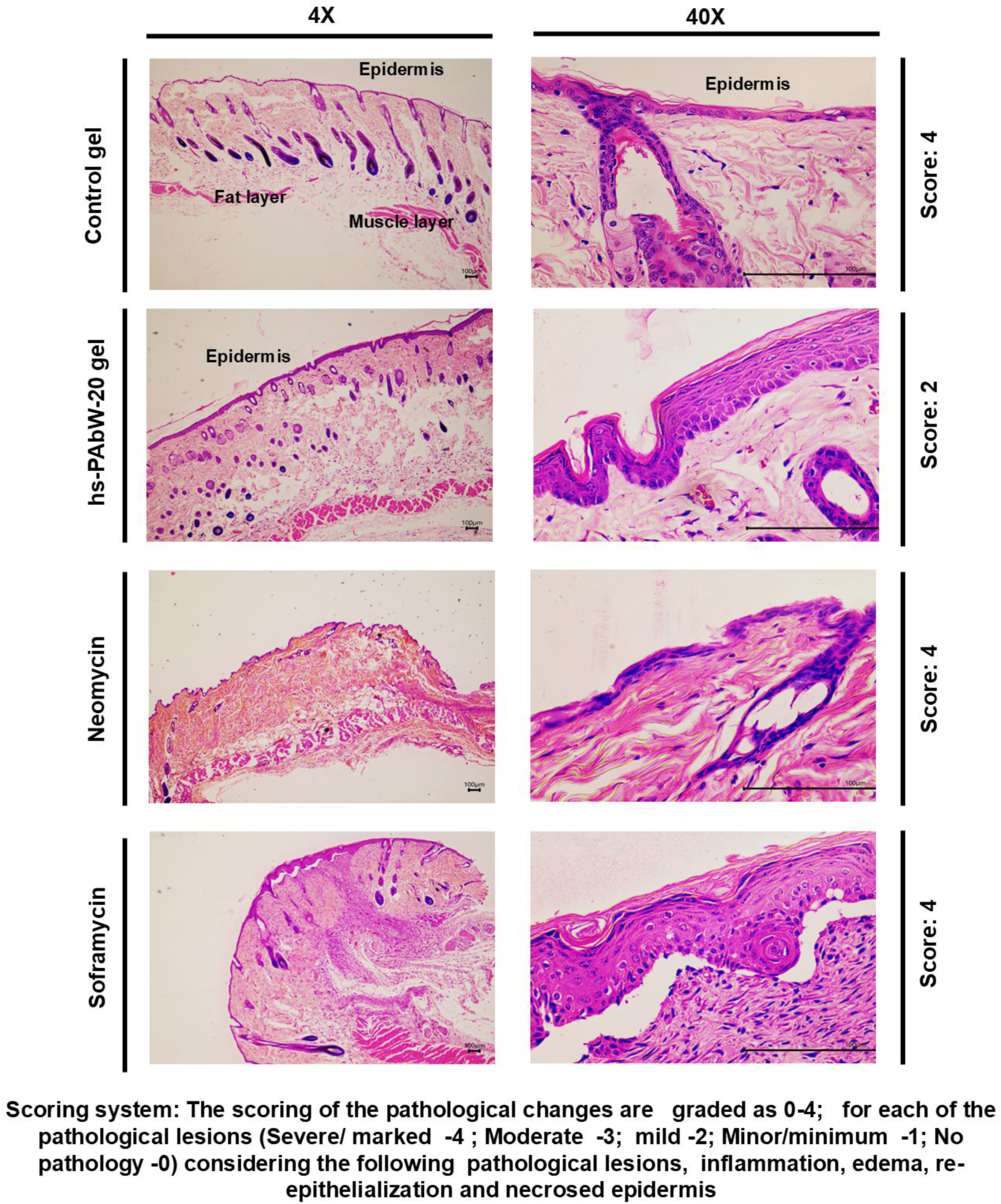
Histopathology of *P. aeruginosa*-infected diabetic wounds at 9d. H&E-stained sections showing tissue repair and epithelialization in wounds under treatment conditions.

Thus, hs-PAbW-20-gel effectively clears infection and enhances healing in diabetic wounds, outperforming standard antibiotic creams.

## Discussion

Wound healing is a tightly orchestrated process disrupted by microbial colonisation, particularly in chronic and diabetic wounds (37–39). Rising antimicrobial resistance has rendered many conventional dressings ineffective, driving the search for novel therapeutics (44, 45).

Here, we present hs-PAbW-20-gel as a clinically viable alternative. hs-PAbW relies on RNS-driven peroxynitrite and nitrotyrosine formation for antimicrobial activity (17–20). Formulated with xanthan gum, the hs-PAbW-20-gel retained full bactericidal potency against *P. aeruginosa* and *S. aureus in vitro* without impairing L929 fibroblast migration.

*In vivo*, alternate-day application of hs-PAbW-20-gel dramatically reduced bacterial burden in infected wounds of both Balb/c and C57BL/6 mice, outperforming or equalling neomycin and soframycin while significantly upregulating key healing genes. Accelerated wound closure was pronounced in Balb/c mice, consistent with EGFR-mediated proliferation and re-epithelialisation. The attenuated response in proinflammatory C57BL/6 mice highlights strain-specific immune modulation (32, 53–56).

Remarkably, in streptozotocin-induced diabetic mice, hs-PAbW-20-gel cleared *P. aeruginosa* more effectively than commercial antibiotics, strongly induced healing markers, and achieved superior histopathological repair—demonstrating particular promise for impaired diabetic wounds (35, 57–59).

To our knowledge, this is the first report of a neutral-pH plasma-activated hydrogel applied successfully *in vivo*. hs-PAbW-20-gel combines potent antimicrobial action with active promotion of tissue repair, positioning it as a next-generation therapeutic for infected and chronic wounds.

## Contributors

Conceptualization: DM, PG, LNR, and DC, Methodology: DM, PG, RSR, KV, AG, Investigation: DM, PG, RSR, KV, AG, Visualization: DM, PG, Data curation: DM, PG, KV, AG, Formal analysis: DM, PG, KV, AG, Fund acquisition: LNR and DC, Project administration: DM, PG, KV, LNR, and DC, Writing-original draft: DM, PG, Editing and proofreading: DM, PG, LNR, and DC, Supervision: LNR and DC.

## Acknowledgments

The authors acknowledge Central Flow Cytometry Facility at IISc.

## Data sharing statement

Authors approve data sharing.

---

**Figure S1.**
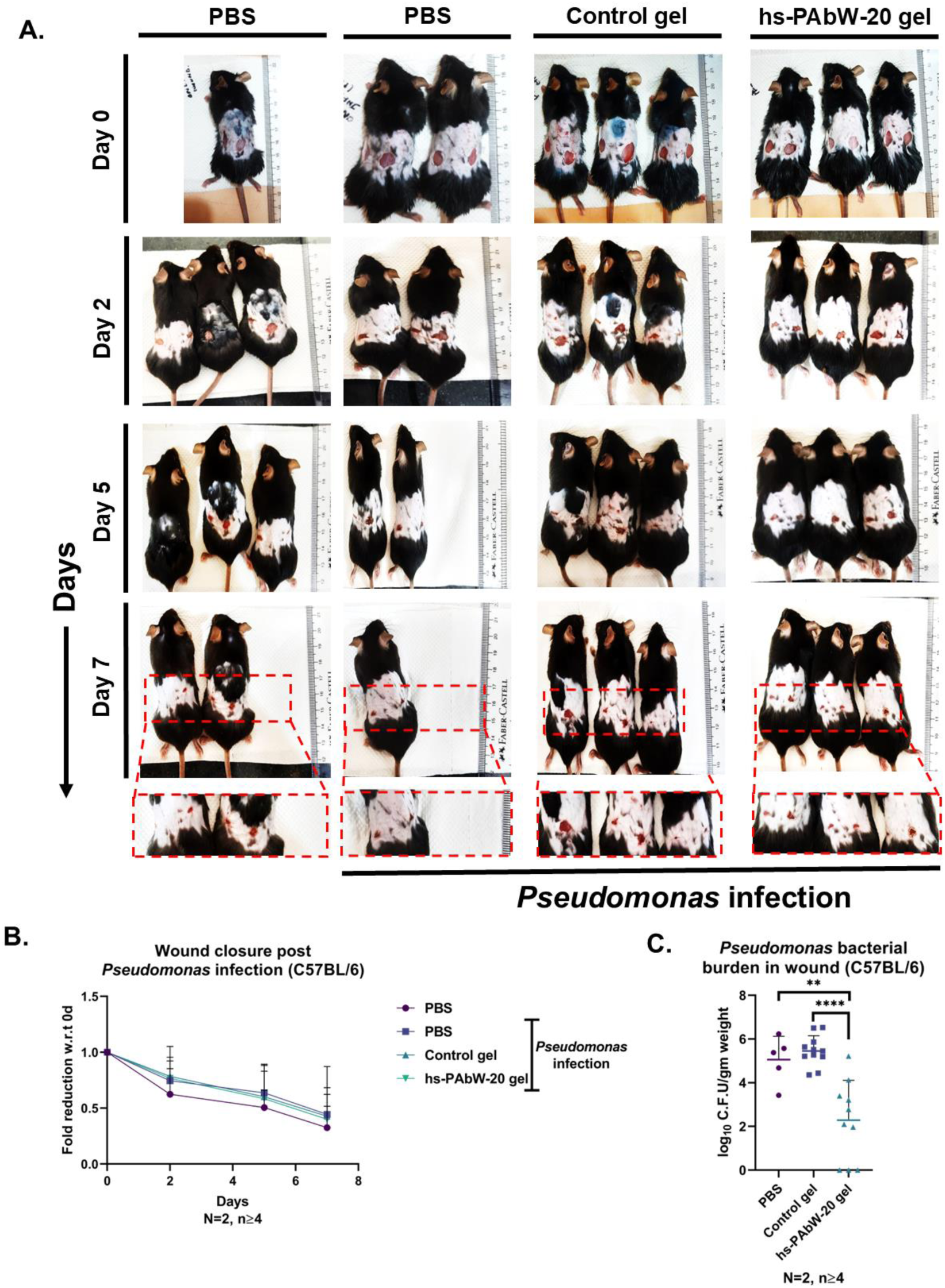
hs-PAbW-20-gel clears *P. aeruginosa* infection in C57BL/6 mice without significantly accelerating macroscopic wound closure. **(A)** Representative wound images (0d–7d) after *P. aeruginosa* infection and treatment with PBS, control gel, or hs-PAbW-20-gel. **(B)** Fold reduction in wound diameter relative to 0d. **(C)** *P. aeruginosa* burden in tissue at 9d. Data in (B–C) are mean±SD, pooled from N=2 independent experiments, n≥4 mice/group. Statistical tests: two-way ANOVA (B), one-way ANOVA (C, grouped), Mann–Whitney U-test (CFU); ****p<0.0001, ***p<0.001, **p<0.01, *p<0.05.

**Figure S2.**
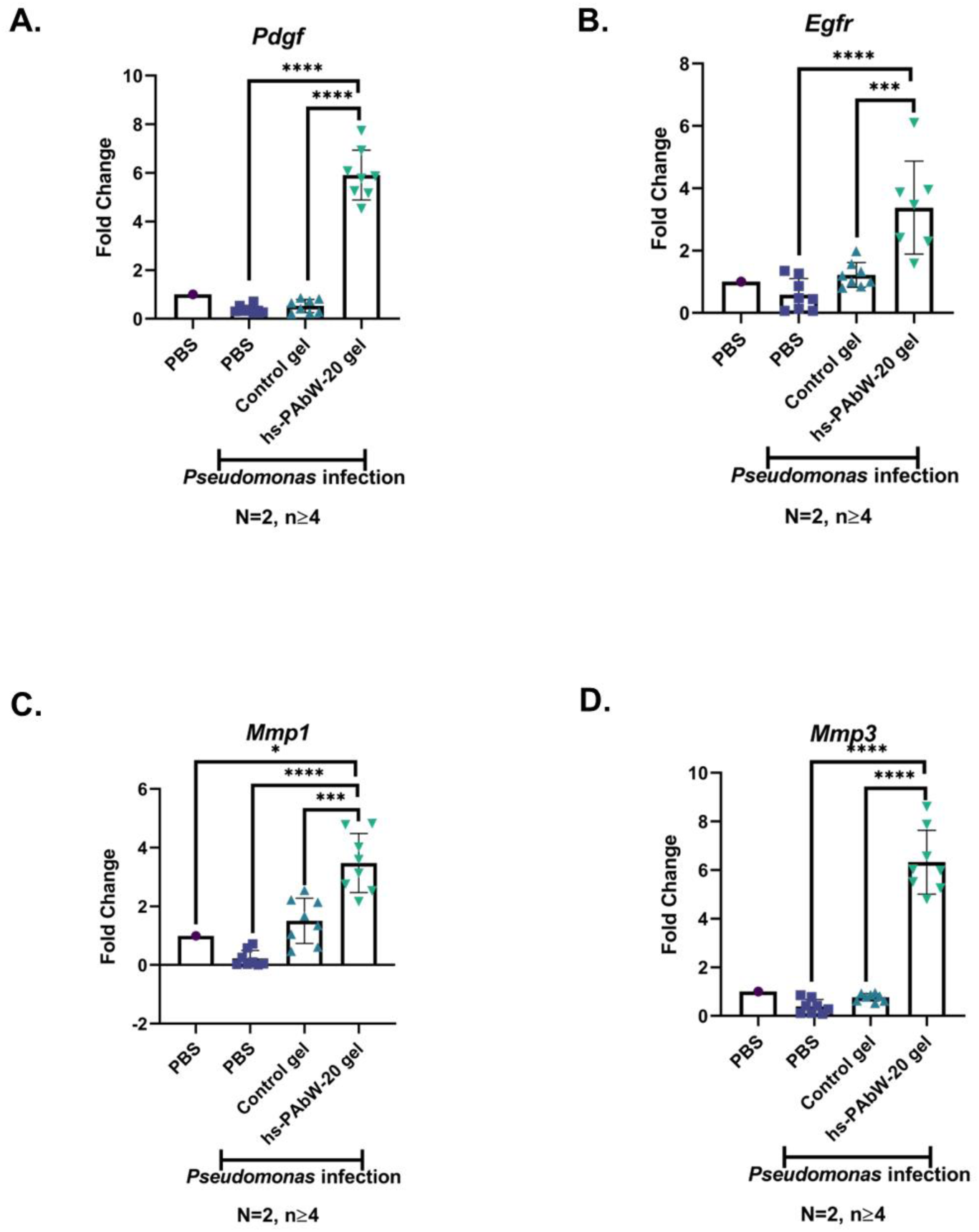
hs-PAbW-20-gel upregulates wound-healing gene expression in *P. aeruginosa*-infected C57BL/6 mouse wounds. Relative mRNA levels of (A) *Pdgf*, (B) *Egfr*, (C) *Mmp1*, and (D) *Mmp3* in uninfected wounds and infected wounds treated with PBS, control gel, or hs-PAbW-20 gel. Data are mean±SD from N=2 independent experiments, n≥4 mice/group. One-way ANOVA; ****p<0.0001, ***p<0.001, **p<0.01, *p<0.05.

**Figure S3.**
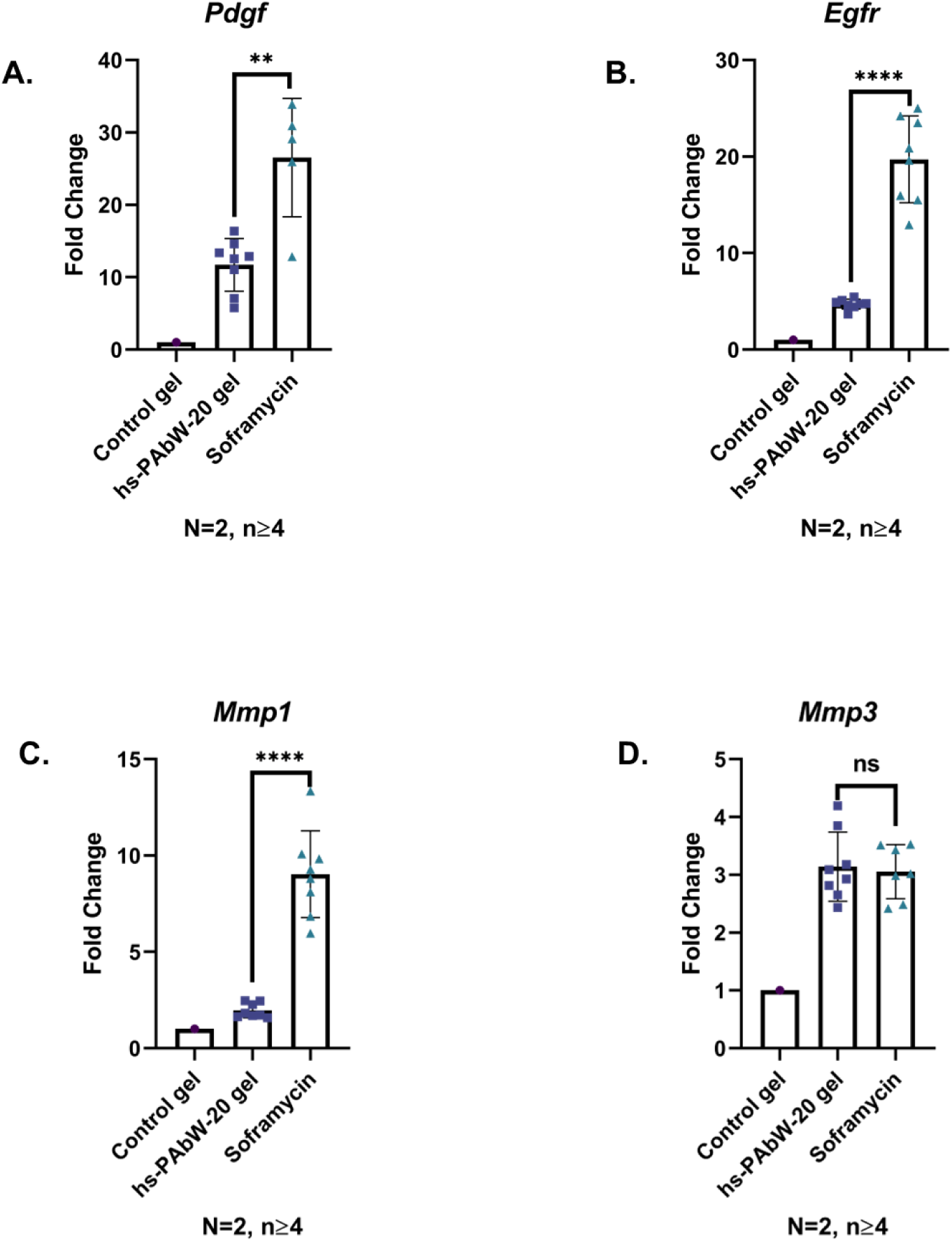
hs-PAbW-20-gel upregulates wound-healing gene expression in *P. aeruginosa*-infected Balb/c mouse wounds, comparable to soframycin. Relative mRNA levels of (A) *Pdgf*, (B) *Egfr*, (C) *Mmp1*, and (D) *Mmp3* in infected wounds treated with control gel, hs-PAbW-20 gel, or soframycin. Data are mean±SD from N=2 independent experiments, n≥4 mice/group. One-way ANOVA; ****p<0.0001, ***p<0.001, **p<0.01, *p<0.05.

**Figure S4.**
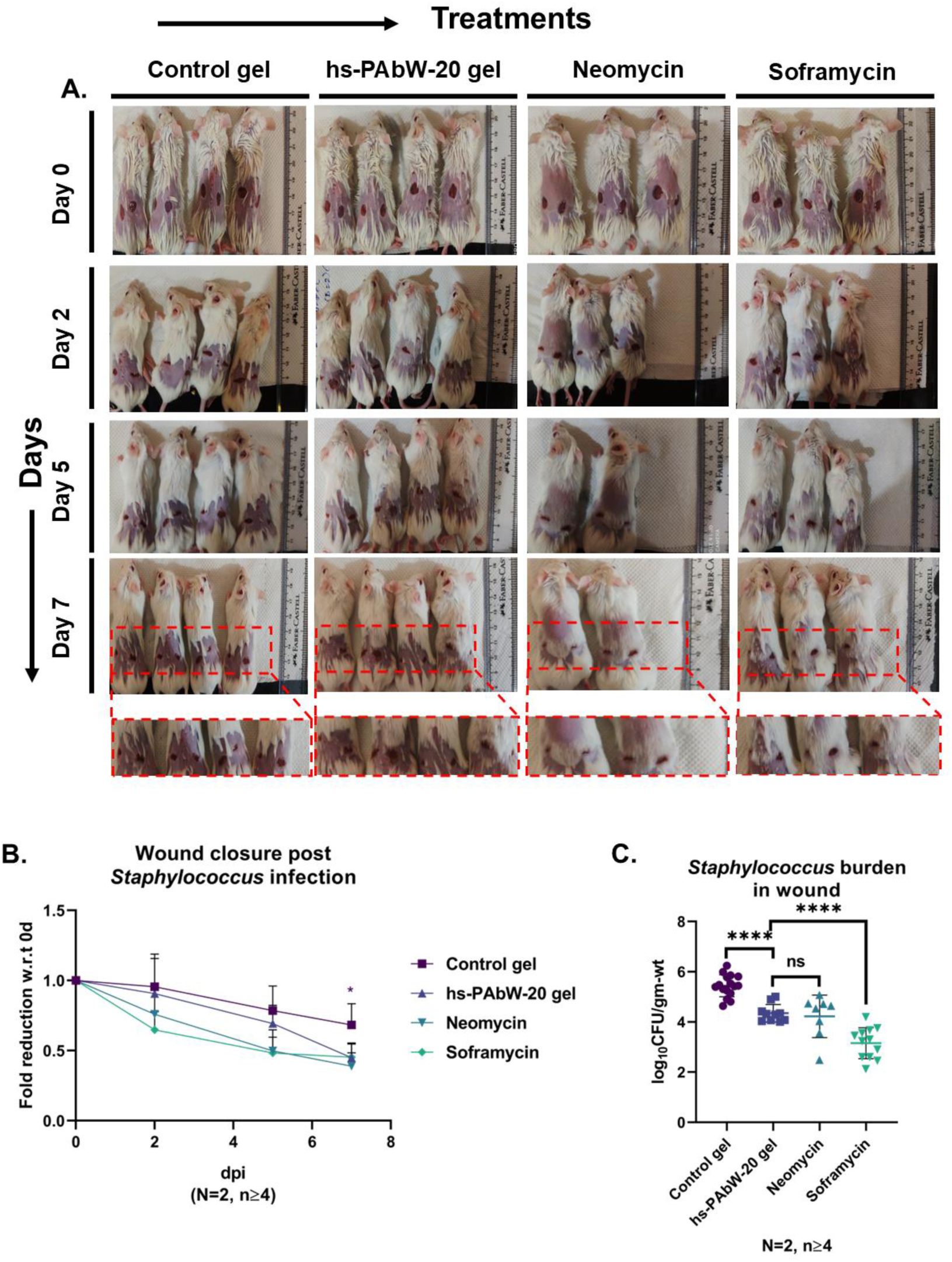
hs-PAbW-20-gel matches commercial antibiotic creams in reducing *S. aureus* burden and accelerating closure in Balb/c mouse wounds. **(A)** Representative wound images (0d–7d) after *S. aureus* infection and treatment with control gel, hs-PAbW-20-gel, neomycin, or soframycin. **(B)** Fold reduction in wound diameter relative to 0d. **(C)** *S. aureus* burden in tissue at 9d. Data in (B–C) are mean±SD, pooled from N=2 independent experiments, n≥4 mice/group. Statistical tests: two-way ANOVA (B), one-way ANOVA (C, grouped), Mann–Whitney U-test (CFU); ****p<0.0001, ***p<0.001, **p<0.01, *p<0.05.

**Figure S5.**
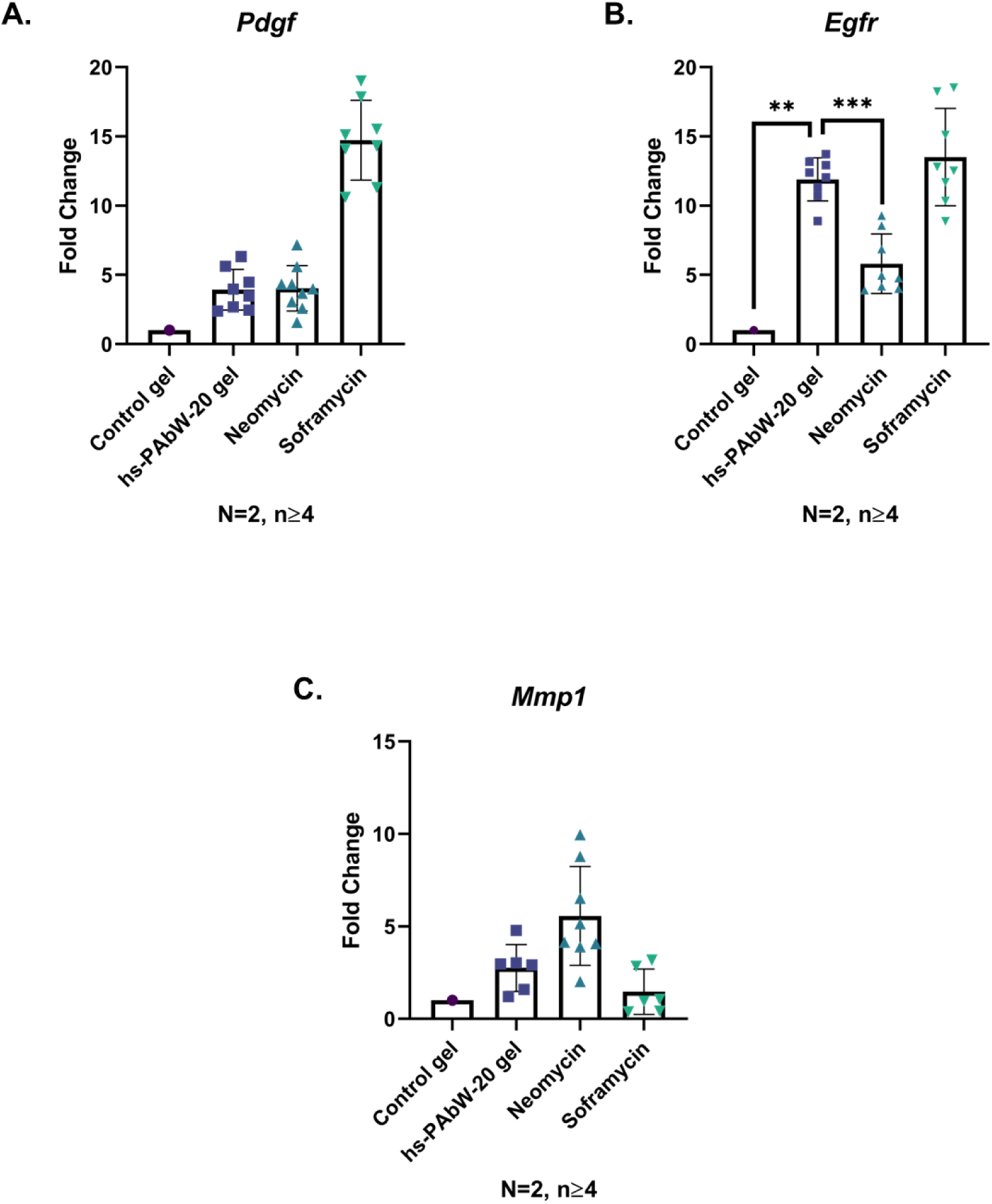
hs-PAbW-20-gel upregulates wound-healing gene expression in *S. aureus*-infected Balb/c mouse wounds. Relative mRNA levels of (A) *Pdgf*, (B) *Egfr*, and (C) *Mmp1* in infected wounds treated with control gel, hs-PAbW-20-gel, neomycin, or soframycin. Data are mean±SD from N=2 independent experiments, n≥4 mice/group. One-way ANOVA; ****p<0.0001, ***p<0.001, **p<0.01, *p<0.05.

**Figure S6.**
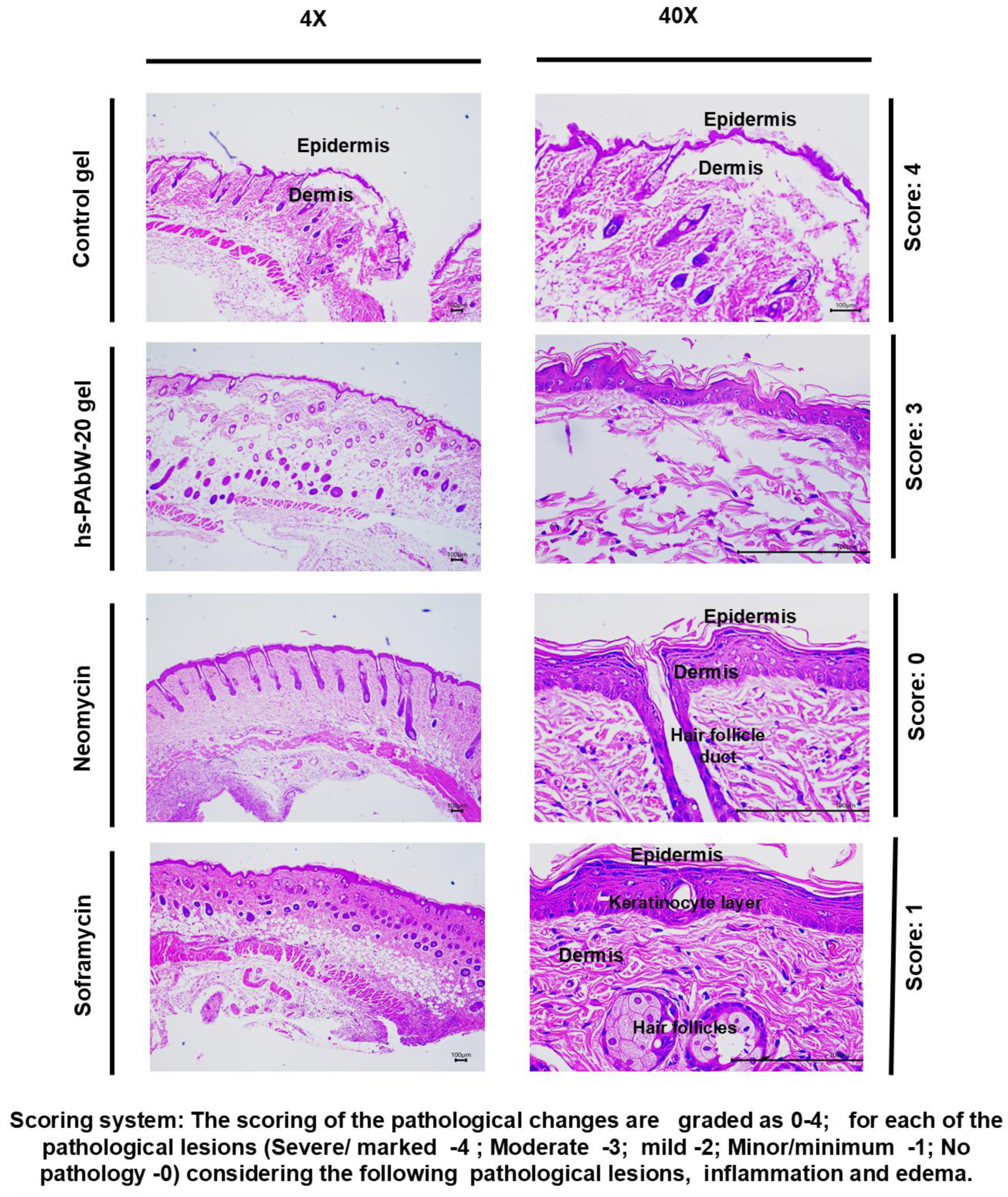
Histopathology of *S. aureus*-infected wounds at 9d. H&E-stained sections showing tissue repair and epithelialization in wounds under treatment conditions.

**Figure S7.**
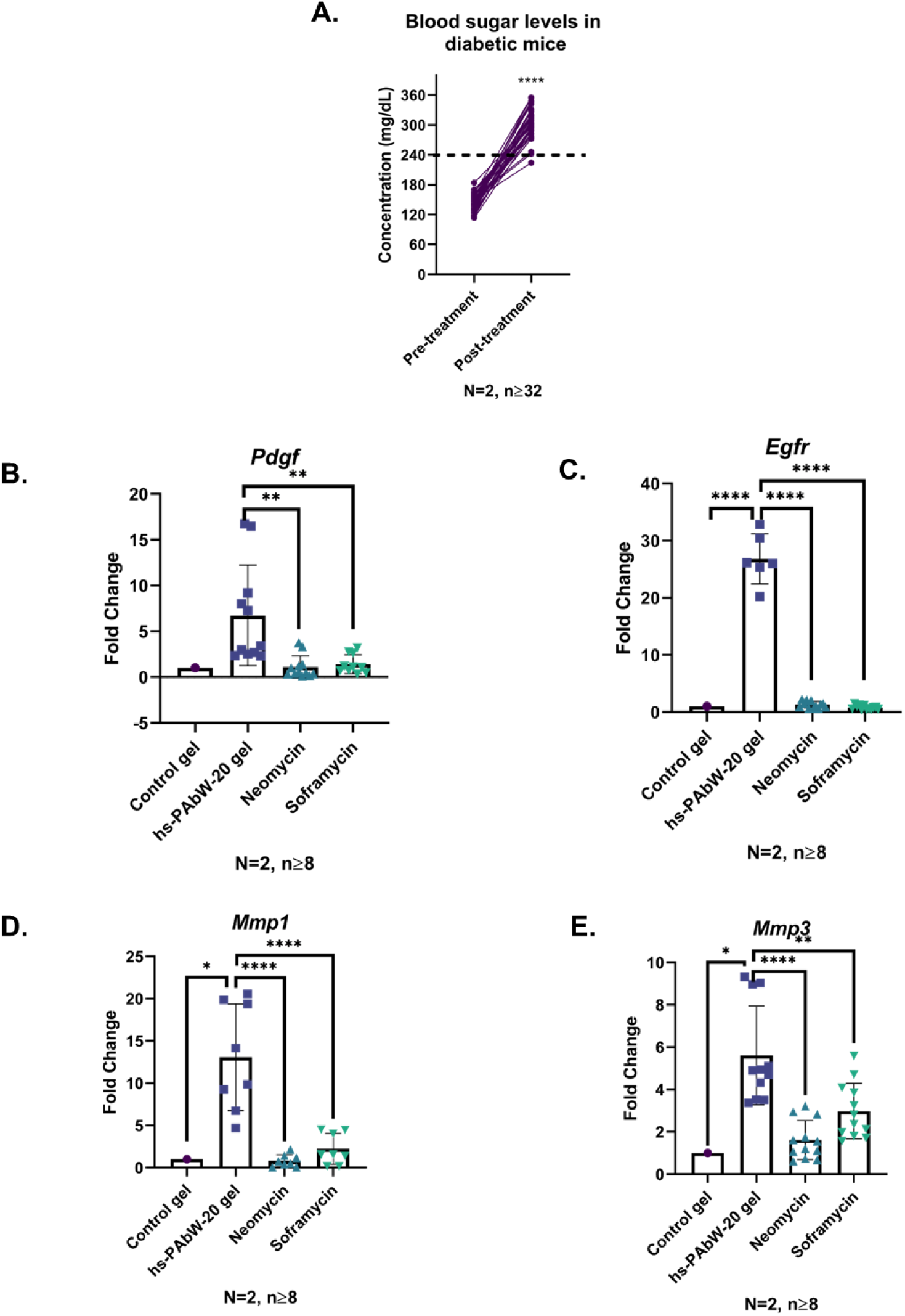
Confirmation of diabetes induction and upregulation of wound-healing genes by hs-PAbW-20-gel in diabetic C57BL/6 mice. **(A)** Blood glucose levels before and after streptozotocin induction (mean±SD; N=2, n≥32). **(B–E)** Relative mRNA expression of (B) *Pdgf*, (C) *Egfr*, (D) *Mmp1*, and (E) *Mmp3* in *P. aeruginosa*-infected diabetic wounds treated with control gel, hs-PAbW-20-gel, neomycin, or soframycin. Data in (B–E) are mean±SD from N=2 independent experiments, n≥8 mice/group. One-way ANOVA; ****p<0.0001, ***p<0.001, **p<0.01, *p<0.05.

## References

1. Arora A, Bharadwaj P, Chaturvedi H, Chowbey P, Gupta S, Leaper D, et al. A review of prevention of surgical site infections in Indian hospitals based on global guidelines for the prevention of surgical site infection, 2016. Journal of Patient Safety and Infection Control. 2018;6(1):1–12.

2. Owens CD, Stoessel K. Surgical site infections: epidemiology, microbiology and prevention. J Hosp Infect. 2008;70 Suppl 2:3–10.

3. Mangram AJ, Horan TC, Pearson ML, Silver LC, Jarvis WR, Committee HICPA. Guideline for prevention of surgical site infection, 1999. Infection Control & Hospital Epidemiology. 1999;20(4):247–80.

4. Emori TG, Gaynes RP. An overview of nosocomial infections, including the role of the microbiology laboratory. Clinical microbiology reviews. 1993;6(4):428–42.

5. Negut I, Grumezescu V, Grumezescu AM. Treatment Strategies for Infected Wounds. Molecules. 2018;23(9).

6. Taati Moghadam M, Khoshbayan A, Chegini Z, Farahani I, Shariati A. Bacteriophages, a New Therapeutic Solution for Inhibiting Multidrug-Resistant Bacteria Causing Wound Infection: Lesson from Animal Models and Clinical Trials. Drug Des Devel Ther. 2020;14:1867–83.

7. Glik J, Kawecki M, Gazdzik T, Nowak M. The impact of the types of microorganisms isolated from blood and wounds on the results of treatment in burn patients with sepsis. Pol Przegl Chir. 2012;84(1):6–16.

8. Maillard JY, Kampf G, Cooper R. Antimicrobial stewardship of antiseptics that are pertinent to wounds: the need for a united approach. JAC Antimicrob Resist. 2021;3(1):dlab027.

9. Eeshita D, Tejashree Anantharaj U, Krishna Karthik M. Antimicrobial Susceptibility Profile of Methicillin Resistant Staphylococcus Aureus (MRSA) Isolates in a Tertiary Care Hospital, Mysuru, India. Indian Journal of Public Health Research & Development. 2023;14(2):88–93.

10. Dash M, Padhi S, Narasimham M, Pattnaik S. Antimicrobial resistance pattern of *Pseudomonas aeruginosa* isolated from various clinical samples in a tertiary care hospital, South Odisha, India. Saudi Journal for Health Sciences. 2014;3(1):15–9.

11. Rajan A, Boopathy B, Radhakrishnan M, Rao L, Schlüter OK, Tiwari BK. Plasma processing: a sustainable technology in agri-food processing. Sustainable Food Technology. 2023;1(1):9–49.

12. Thirumdas R, Kothakota A, Annapure U, Siliveru K, Blundell R, Gatt R, et al. Plasma activated water (PAW): Chemistry, physico-chemical properties, applications in food and agriculture. Trends in Food Science & Technology. 2018;77:21–31.

13. Girard PM, Arbabian A, Fleury M, Bauville G, Puech V, Dutreix M, et al. Synergistic Effect of H2O2 and NO2 in Cell Death Induced by Cold Atmospheric He Plasma. Sci Rep. 2016;6:29098.

14. Hoeben WFLM, van Ooij PP, Schram DC, Huiskamp T, Pemen AJM, Lukeš P. On the Possibilities of Straightforward Characterization of Plasma Activated Water. Plasma Chemistry and Plasma Processing. 2019;39(3):597–626.

15. Punith N, Harsha R, Lakshminarayana R, Hemanth M, S. Anand M, Dasappa S. Plasma Activated Water Generation and its Application in Agriculture. Advanced Materials Letters. 2019;10(10):700–4.

16. Han L, Boehm D, Patil S, Cullen PJ, Bourke P. Assessing stress responses to atmospheric cold plasma exposure using Escherichia coli knock-out mutants. J Appl Microbiol. 2016;121(2):352–63.

17. N. P, Singh AK, J. A, Boopathy B, Chatterjee R, M. H, et al. Generation of neutral pH high-strength plasma-activated water from a pin to water discharge and its bactericidal activity on multidrug-resistant pathogens. Plasma Processes and Polymers. 2023;20(1):2200133.

18. Zhou R, Zhou R, Prasad K, Fang Z, Speight R, Bazaka K, et al. Cold atmospheric plasma activated water as a prospective disinfectant: the crucial role of peroxynitrite. Green Chemistry. 2018;20(23):5276–84.

19. Boopathy B, Mukherjee D, Nishanth V, Chowdhury AR, Chakravortty D, Rao L. Generation of Species-Specific High-Strength Plasma Activated Water at Neutral pH and its Antimicrobial Characteristics. Plasma Chemistry and Plasma Processing. 2024;44(2):1003–17.

20. Mukherjee D, Roy Chowdhury A, Ghosh P, Vishwa N, Rao L, Chakravortty D. Reactive nitrogen species (RNS) and its reaction intermediates with reactive oxygen species (ROS) in pH-neutral high-strength plasma-activated water determines the antimicrobial activity against ESKAPE Pathogens. bioRxiv. 2024:2024.10. 06.616848.

21. Chowdhury AR, Mukherjee D, Singh AK, Chakravortty D. Loss of outer membrane protein A (OmpA) impairs the survival of Salmonella Typhimurium by inducing membrane damage in the presence of ceftazidime and meropenem. J Antimicrob Chemother. 2022;77(12):3376–89.

22. Mukherjee D, Wagle S, Ghosh P, Biswas A, Chakravortty D. Intracellular formate is a critical mediator of the antimicrobial fate of Salmonella Typhimurium Against meropenem and ciprofloxacin. 2025.

23. Bolla SR, Mohammed Al-Subaie A, Yousuf Al-Jindan R, Papayya Balakrishna J, Kanchi Ravi P, Veeraraghavan VP, et al. In vitro wound healing potency of methanolic leaf extract of Aristolochia saccata is possibly mediated by its stimulatory effect on collagen-1 expression. Heliyon. 2019;5(5):e01648.

24. Liang CC, Park AY, Guan JL. In vitro scratch assay: a convenient and inexpensive method for analysis of cell migration in vitro. Nat Protoc. 2007;2(2):329–33.

25. Huang Y, Yang N, Teng D, Mao R, Hao Y, Ma X, et al. Antibacterial peptide NZ2114-loaded hydrogel accelerates Staphylococcus aureus-infected wound healing. Appl Microbiol Biotechnol. 2022;106(9-10):3639–56.

26. Furman BL. Streptozotocin-Induced Diabetic Models in Mice and Rats. Curr Protoc. 2021;1(4):e78.

27. Serra R, Grande R, Butrico L, Rossi A, Settimio UF, Caroleo B, et al. Chronic wound infections: the role of Pseudomonas aeruginosa and Staphylococcus aureus. Expert Rev Anti Infect Ther. 2015;13(5):605–13.

28. Alonso-Matilla R, Provenzano PP, Odde DJ. Physical principles and mechanisms of cell migration. NPJ Biol Phys Mech. 2025;2(1):2.

29. Barrientos S, Stojadinovic O, Golinko MS, Brem H, Tomic-Canic M. Growth factors and cytokines in wound healing. Wound Repair Regen. 2008;16(5):585–601.

30. Armstrong DG, Jude EB. The role of matrix metalloproteinases in wound healing. J Am Podiatr Med Assoc. 2002;92(1):12–8.

31. Kandhwal M, Behl T, Singh S, Sharma N, Arora S, Bhatia S, et al. Role of matrix metalloproteinase in wound healing. Am J Transl Res. 2022;14(7):4391–405.

32. Fukushima A, Yamaguchi T, Ishida W, Fukata K, Taniguchi T, Liu FT, et al. Genetic background determines susceptibility to experimental immune-mediated blepharoconjunctivitis: comparison of Balb/c and C57BL/6 mice. Exp Eye Res. 2006;82(2):210–8.

33. Wang Y, Zhang Y, Yang YP, Jin MY, Huang S, Zhuang ZM, et al. Versatile dopamine-functionalized hyaluronic acid-recombinant human collagen hydrogel promoting diabetic wound healing via inflammation control and vascularization tissue regeneration. Bioact Mater. 2024;35:330–45.

34. Liu W, Gao R, Yang C, Feng Z, Ou-Yang W, Pan X, et al. ECM-mimetic immunomodulatory hydrogel for methicillin-resistant Staphylococcus aureus-infected chronic skin wound healing. Sci Adv. 2022;8(27):eabn7006.

35. He J, Chen J, Liu T, Qin F, Wei W. Research Progress of Multifunctional Hydrogels in Promoting Wound Healing of Diabetes. Int J Nanomedicine. 2025;20:7549–78.

36. Rubinstein MR, Genaro AM, Wald MR. Differential effect of hyperglycaemia on the immune response in an experimental model of diabetes in BALB/cByJ and C57Bl/6J mice: participation of oxidative stress. Clin Exp Immunol. 2013;171(3):319–29.

37. Huang C, Dong L, Zhao B, Lu Y, Huang S, Yuan Z, et al. Anti-inflammatory hydrogel dressings and skin wound healing. Clinical and translational medicine. 2022;12(11):e1094.

38. Dąbrowska A, Spano F, Derler S, Adlhart C, Spencer ND, Rossi RM. The relationship between skin function, barrier properties, and body-dependent factors. Skin Research and Technology. 2018;24(2):165–74.

39. Hu P, Yang Q, Wang Q, Shi C, Wang D, Armato U, et al. Mesenchymal stromal cells-exosomes: a promising cell-free therapeutic tool for wound healing and cutaneous regeneration. Burns & trauma. 2019;7.

40. Rahimnejad M, Derakhshanfar S, Zhong W. Biomaterials and tissue engineering for scar management in wound care. Burns & trauma. 2017;5.

41. Murray RZ, West ZE, Cowin AJ, Farrugia BL. Development and use of biomaterials as wound healing therapies. Burns & trauma. 2019;7.

42. Tehrany PM, Rahmanian P, Rezaee A, Ranjbarpazuki G, Sohrabi Fard F, Asadollah Salmanpour Y, et al. Multifunctional and theranostic hydrogels for wound healing acceleration: An emphasis on diabetic-related chronic wounds. Environ Res. 2023;238(Pt 1):117087.

43. Bowler PG. Wound pathophysiology, infection and therapeutic options. Ann Med. 2002;34(6):419–27.

44. Zaman SB, Hussain MA, Nye R, Mehta V, Mamun KT, Hossain N. A Review on Antibiotic Resistance: Alarm Bells are Ringing. Cureus. 2017;9(6):e1403.

45. Cristea Hohota AG, Lisa EL, Iacob Ciobotaru S, Dragostin I, Stefan CS, Fulga I, et al. Antimicrobial Smart Dressings for Combating Antibiotic Resistance in Wound Care. Pharmaceuticals (Basel). 2025;18(6).

46. Villalba-Rodríguez AM, Martínez-González S, Sosa-Hernández JE, Parra-Saldívar R, Bilal M, Iqbal HM. Nanoclay/polymer-based hydrogels and enzyme-loaded nanostructures for wound healing applications. Gels. 2021;7(2):59.

47. Yao H, Wu M, Lin L, Wu Z, Bae M, Park S, et al. Design strategies for adhesive hydrogels with natural antibacterial agents as wound dressings: Status and trends. Materials Today Bio. 2022;16:100429.

48. Herianto S, Hou CY, Lin CM, Chen HL. Nonthermal plasma-activated water: A comprehensive review of this new tool for enhanced food safety and quality. Comprehensive Reviews in Food Science and Food Safety. 2021;20(1):583–626.

49. Rathore V, Nema SK. Optimization of process parameters to generate plasma activated water and study of physicochemical properties of plasma activated solutions at optimum condition. Journal of Applied Physics. 2021;129(8).

50. Hoeben W, Van Ooij P, Schram D, Huiskamp T, Pemen A, Lukeš P. On the possibilities of straightforward characterization of plasma activated water. Plasma Chemistry and Plasma Processing. 2019;39(3):597–626.

51. Punith N, Harsha R, Lakshminarayana R, Hemanth M, S Anand M, Dasappa S. Plasma activated water generation and its application in agriculture. Advanced Materials Letters. 2019;10(10):700–4.

52. Machala Z, Tarabová B, Sersenová D, Janda M, Hensel K. Chemical and antibacterial effects of plasma activated water: Correlation with gaseous and aqueous reactive oxygen and nitrogen species, plasma sources and air flow conditions. Journal of Physics D: Applied Physics. 2018;52(3):034002.

53. Mao Y, Ma J, Xia Y, Xie X. The Overexpression of Epidermal Growth Factor (EGF) in HaCaT Cells Promotes the Proliferation, Migration, Invasion and Transdifferentiation to Epidermal Stem Cell Immunophenotyping of Adipose-Derived Stem Cells (ADSCs). Int J Stem Cells. 2020;13(1):93–103.

54. Saaristo A, Tammela T, Farkkila A, Karkkainen M, Suominen E, Yla-Herttuala S, et al. Vascular endothelial growth factor-C accelerates diabetic wound healing. Am J Pathol. 2006;169(3):1080–7.

55. Teller P, White TK. The physiology of wound healing: injury through maturation. Surg Clin North Am. 2009;89(3):599–610.

56. Merkulova Y, Shen Y, Parkinson LG, Raithatha SA, Zhao H, Westendorf K, et al. Granzyme B inhibits keratinocyte migration by disrupting epidermal growth factor receptor (EGFR)-mediated signaling. Biol Chem. 2016;397(9):883–95.

57. Qi X, Cai E, Xiang Y, Zhang C, Ge X, Wang J, et al. An Immunomodulatory Hydrogel by Hyperthermia-Assisted Self-Cascade Glucose Depletion and ROS Scavenging for Diabetic Foot Ulcer Wound Therapeutics. Adv Mater. 2023;35(48):e2306632.

58. Cao J, Wu B, Yuan P, Liu Y, Hu C. Rational Design of Multifunctional Hydrogels for Wound Repair. J Funct Biomater. 2023;14(11).

59. Peng Y, Guo Y, Ge X, Gong Y, Wang Y, Ou Z, et al. Construction of programmed time-released multifunctional hydrogel with antibacterial and anti-inflammatory properties for impaired wound healing. J Nanobiotechnology. 2024;22(1):126.

